# K27-linked ubiquitination restrains the NOD2-RIP2 signaling pathway

**DOI:** 10.64898/2026.01.16.699886

**Authors:** Swarupa Panda, Amitha Kamath, Nelson O. Gekara

**Author notes:** Correspondence to: **Nelson O. Gekara** and **Swarupa Panda**. These authors share correspondence responsibility (Swarupa Panda, Nelson O. Gekara).

## Abstract

Nucleotide-binding oligomerization domain 2 (NOD2) signaling is a critical arm of host defense and immune homeostasis but requires strict regulation to prevent self-injury. Central to this regulation is the polyubiquitination of the adaptor protein RIP2, yet the precise ubiquitin linkages and their dynamic control remain incompletely understood. Here, we identify K27-linked polyubiquitination of RIP2 by the E3 ligase XIAP as a novel inhibitory signal that attenuates RIP2 complex assembly and downstream NF-κB activation, thereby fine-tuning NOD2-induced inflammatory signaling. We show that XIAP-mediated K63-linked ubiquitination promotes NOD2-RIP2 activation, whereas K27-linked ubiquitination impairs the recruitment of TAK1 and NEMO to the NOD2-RIP2 complex, dampening downstream signaling and cytokine production. We further demonstrate that this regulation is counterbalanced by the deubiquitinase MYSM1, which selectively removes K27-, K63-. and M1-polyubiquitin chains via its SWIRM-MPN domains to restore balanced signaling. Together, these findings define a dynamic XIAP-RIP2-MYSM1 axis in which opposing ubiquitin linkages fine-tune NOD2 responses, establishing a molecular rheostat for immune signaling with implications for inflammatory disease pathogenesis and opportunities for targeted therapeutic intervention.

## Introduction

An optimal host defense requires a finely balanced regulation of pattern recognition receptors (PRRs), as their dysregulation is associated with numerous inflammatory and autoimmune disorders(Akira *et al*, 2006; Takeuchi & Akira, 2010). Among PRRs, nucleotide-binding oligomerization domain 2 (NOD2) is pivotal for sensing bacterial peptidoglycan-derived muramyl dipeptide and activating NF-κB and MAPK signaling to induce inflammatory cytokines and antimicrobial peptides (Girardin *et al*, 2003; Park *et al*, 2007). Its clinical relevance is underscored by associations of NOD2 mutations with Crohn’s disease (in the LRR domain) and Blau syndrome (in the NOD domain). NOD2 is also implicated in allergic inflammation, with increased signaling contributing to asthma and related conditions(Hugot *et al*, 2001; Mao *et al*, 2022; Parackova *et al*, 2020; Zhang *et al*, 2024).

Ubiquitination, a highly conserved post-translational modification, is essential for regulating diverse cellular processes, including innate immune responses (Bhoj & Chen, 2009; Hu & Sun, 2016; Li *et al*, 2016; Liu *et al*, 2016) by modulating protein stability, localization, and signaling complex formation (Akutsu *et al*, 2016; Sheng *et al*, 2024). Ubiquitin is a small 8 kDa protein that can be covalently attached to substrates as a single unit or as polyubiquitin chains linked via any of its seven lysine residues (K6, K11, K27, K29, K33, K48, K63) or its N-terminal methionine (M1) for linear chains(Agrata & Komander, 2025; Akutsu *et al*., 2016; Damgaard, 2021; Yau & Rape, 2016). While K48-linked polyubiquitin chains predominantly target proteins for proteasomal degradation, K63 and M1 linkages are widely recognized for their non-degradative roles in immune signaling, including the formation of signaling scaffolds and regulation of protein-protein interactions(Chau *et al*, 1989; Chen & Sun, 2009; Komander *et al*, 2009; Liu *et al*, 2024; Tokunaga *et al*, 2009). However the function of the non-canonical ubiquitin linkages such as K6, K11, K27, K29, and K33 (Akizuki *et al*, 2024; Harris *et al*, 2021; Jiang *et al*, 2023; Michel *et al*, 2017; Zhao *et al*, 2021) in immune regulation, and the enzymes involved in their assembly and removal is less clear.

Ubiquitination is now a well-established regulator of NOD2 signaling. K63-linked polyubiquitination of RIP2, particularly at lysine 209, is essential for recruiting the TAK1 complex and activating NF-κB (Panda & Gekara, 2018; Pellegrini *et al*, 2018; Windheim *et al*, 2007). E3 ligases including cIAPs, Pellino3, ITCH, and XIAP have been implicated in K63-linked polyubiquitination of RIP2(Bertrand *et al*, 2009; Goncharov *et al*, 2018; Krieg *et al*, 2009; Lu *et al*, 2025; Stafford *et al*, 2018; Tao *et al*, 2009; Yang *et al*, 2013). The linear ubiquitin assembly complex (LUBAC)-comprising RNF31/HOIP adds M1-linked chains to NEMO, further amplifying NF-κB activation (Damgaard *et al*, 2012; Draber *et al*, 2015; Tokunaga *et al*., 2009).

K27-linked polyubiquitination is best known for regulating protein fate, but also contributes to cellular homeostasis by influencing processes such as the DNA damage response and mitochondrial quality control (Castañeda *et al*, 2016; Gatti *et al*, 2015; Jiang *et al*., 2023; Shearer *et al*, 2022). In NOD2 signaling, RIP2 can carry K27-linked chains often as heterotypic oligomers with K63 and M1 linkages (Panda & Gekara, 2018). However, the impact of K27-linked polyubiquitins on NOD2-RIP2 signaling and the identity of the E3 ligase and deubiquitinase responsible for the attachment and removal of these polyubiquitins remains unknown.

Here, we reveal K27-linked polyubiquitination as a key negative regulator of NOD2 signaling that limits RIP2 complex assembly and NF-κB activation. The E3 ligase XIAP deposits these inhibitory K27 chains alongside activating K63 linkages on RIP2, while deubiquitinase MYSM1 counterbalances this by stripping both via its SWIRM–MPN domains. This XIAP-RIP2-MYSM1 axis thus establishes a ubiquitin-based rheostat, and a previously unrecognized regulatory layer for fine-tuning NOD2-driven innate immune responses.

## Results

### Characterization of RIP2-associated polyubiquitinchains

By performing immunoblot analysis of whole-cell lysates from bone marrow-derived macrophages stimulated with the NOD2 agonist L18-MDP for varying durations, we found that NOD2 activation was associated with an increase in K63-, K48-, K27-, and M1-linked polyubiquitin chains, along with enhanced phosphorylation of downstream signaling markers TAK1 (p-TAK1) and RIP2 (p-RIP2) (Fig 1A). A focal point of NOD2 signaling is RIP2, which nucleates the NOD2:RIP2 complex and enables downstream signaling events leading to inflammatory gene expression. Tandem Ubiquitin Binding Entity (TUBE) pull-down assays followed by immunoblotting not only confirmed the presence of K63-, M1-, K48-, and K27-linked ubiquitin chains but also revealed that, following NOD2 activation, RIP2 was polyubiquitinated (Fig1B). To further define these modifications, we immunoprecipitated RIP2 from native or denatured lysates (boiled in SDS to disrupt non-covalent interactions) of L18-MDP-stimulated BMDMs and probed with linkage-specific antibodies. This analysis demonstrated that K63-, K48-, K27-, and M1-linked ubiquitin chains were covalently attached to RIP2 (Fig 1C). Collectively, these results show that NOD2 activation induces covalent attachment of multiple ubiquitin linkage types to RIP2, generating a heterogeneous polyubiquitin landscape that may coordinate and finetune downstream signaling (Fig 1D).

**Figure 1.**
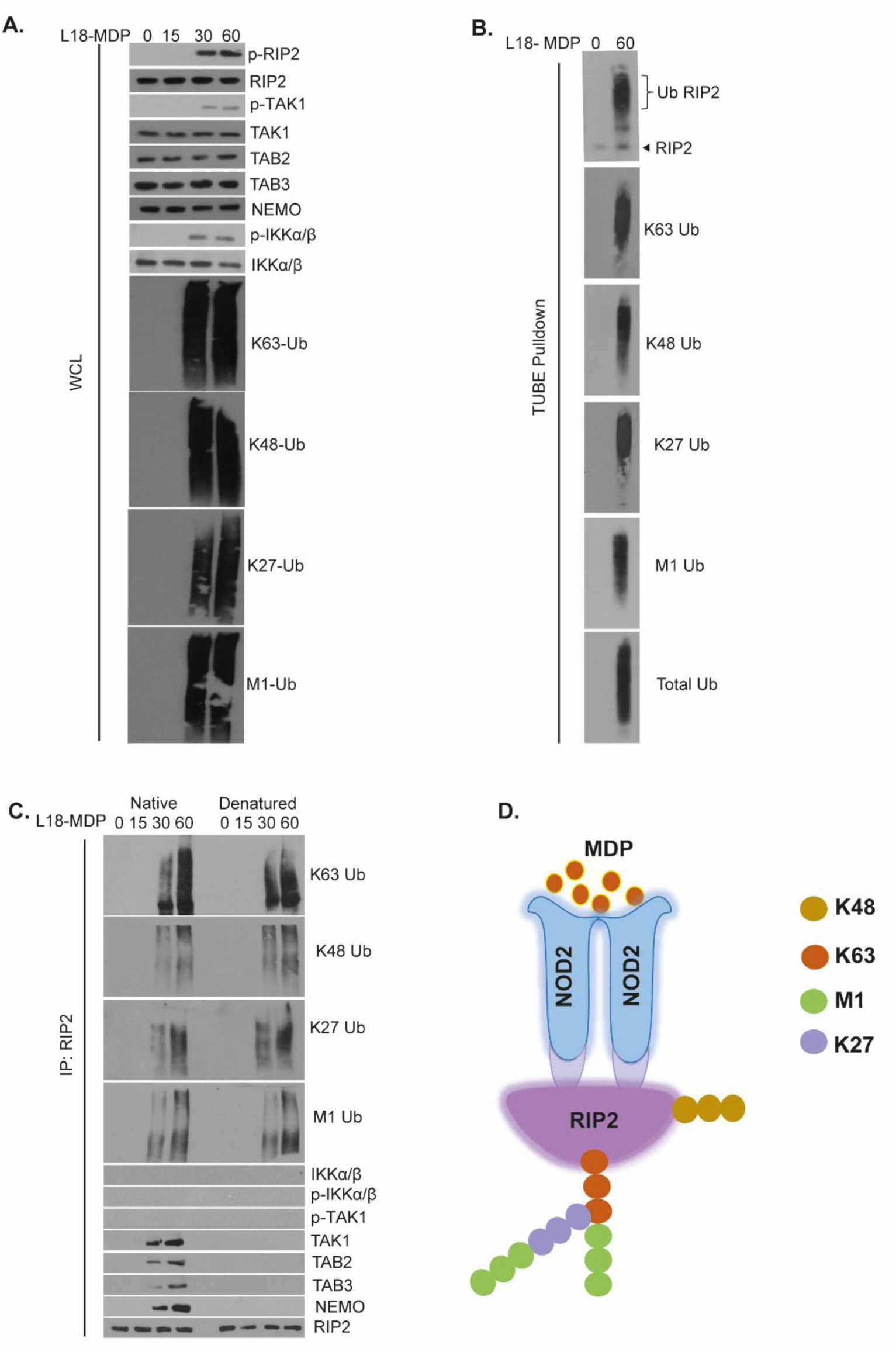
RIP2 is conjugated with multiple polyubiquitin chains NOD2 Activation. **(A)** Whole-cell lysates (WCL) from wild-type (WT) bone marrow-derived macrophages (BMDMs) stimulated with L18-MDP were assessed by immunoblotting for the indicated signaling proteins and polyubiquitin chains. **(B)** Tandem ubiquitin-binding entity (TUBE) pull-down of ubiquitinated proteins from WT BMDMs stimulated with L18-MDP. RIP2-associated ubiquitin linkage types were detected using linkage-specific antibodies. The experiment shown is representative of three independent biological replicates. **(C)** Native and denatured immunoprecipitation (IP) of RIP2 from WT BMDMs stimulated with L18-MDP for the indicated time points were assessed by immunoblotting for the indicated signaling proteins and polyubiquitin chains. **(D)** Schematic summarizing the polyubiquitin linkages identified on RIP2. RIP2 is modified with K48-, K63-, M1-, and K27-linked polyubiquitin chains, each contributing to the regulation of downstream NOD2 signaling pathways. Representative blots from at least three independent experiments are shown.

### K27-linked polyubiquitin chains limit NOD2-induced RIP2 signaling and Impair RIP2 Complex Assembly

K63-and M1-linked polyubiquitin chains have previously been established as essential drivers of NOD2-RIP2 signaling (Hrdinka *et al*, 2016; Panda & Gekara, 2018). However, the impact of K27-linked chains on NOD2 signaling remains unknown. To address this, HEK293 cells were transfected with plasmids expressing NOD2, Flag-RIP2, and HA-tagged ubiquitin mutants restricted to single lysine linkages (K6, K11, K27, K29, K33, K48, K63) or M1 (KO). After L18-MDP stimulation (60 min), ubiquitinated proteins were immunoprecipitated using anti-HA antibodies. Co-immunoprecipitation revealed RIP2 conjugation only with K27-, K48-, K63-, and M1-linked chains; K6, K11, K29, and K33 linkages showed no detectable association (Fig 2A). To assess the functional impact of specific ubiquitin linkages, we measured NF-κB transcriptional activity in HEK293 cells co-transfected with RIP2 and HA-tagged ubiquitin mutants. HA-K63 Ub (forming only K63- and M1-linked chains) enhanced NF-κB-driven luciferase activity, IL-6 secretion, and TNFα production after NOD2 stimulation (Fig 2B and C). In contrast, HA-K27 Ub (forming only K27- and M1-linked chains) suppressed NF-κB activation and reduced IL-6/TNFα levels after MDP treatment, revealing a repressive role for K27 linkages in NOD2-RIP2 signaling. The HA-K63+K27 Ub variant (forming K63-, K27-, and M1-linked chains) failed to restore full activation, indicating K27 dominance (Fig 2B and C).

**Figure 2.**
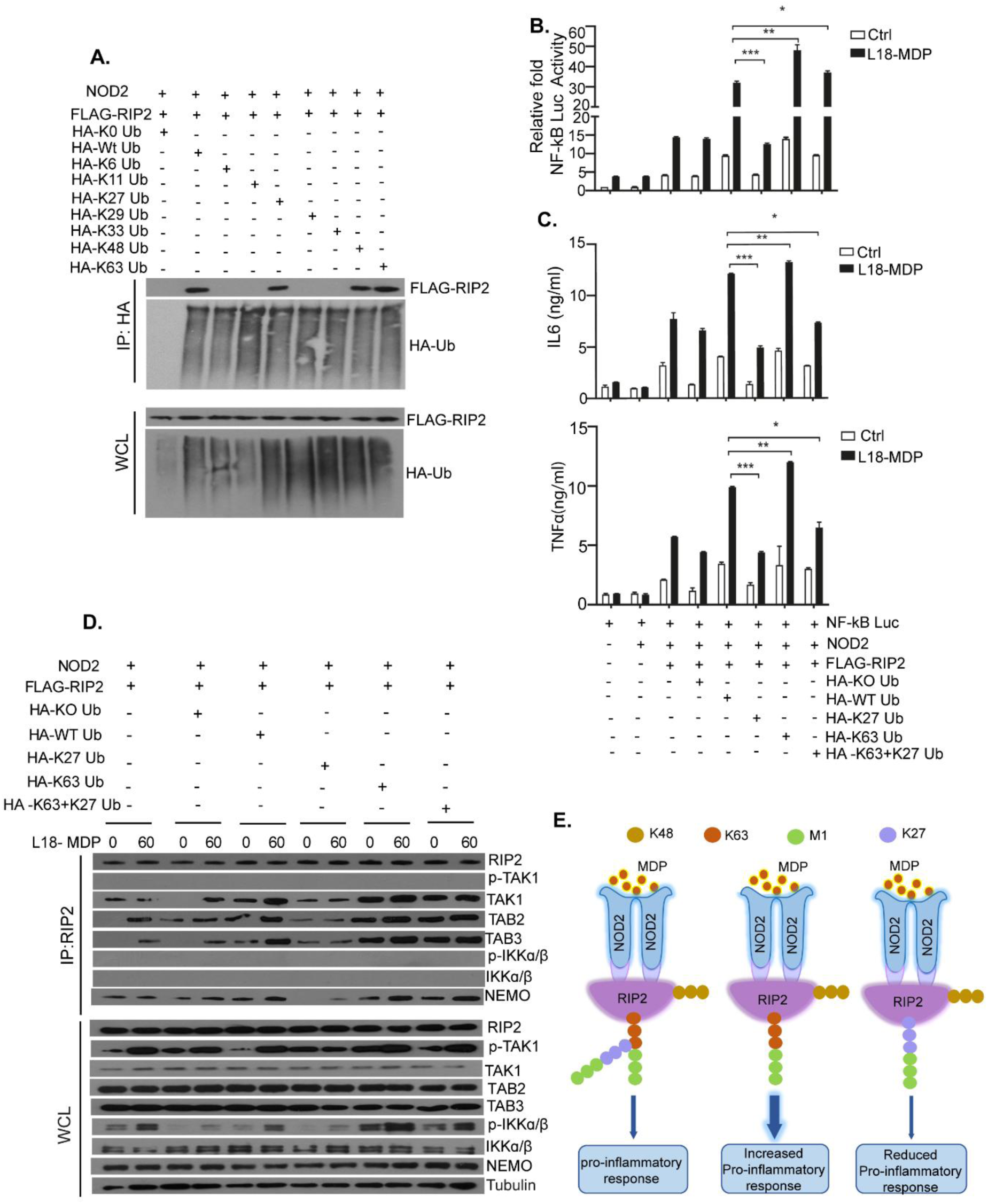
K27-linked polyubiquitin chains suppress NO2-RIP2 signaling by impeding RIP2 complex assembly and downstream cytokine responses. **(A)** HEK293 cells were co-transfected with plasmids encoding NOD2, FLAG-tagged RIP2, and HA-tagged ubiquitin (HA-Ub) variants: wild-type (WT), lysine-less KO (all lysines mutated to arginine), or K-only (single lysine preserved: K6, K11, K27, K33, K48, or K63). Cells were then stimulated with L18-MDP for 60 min, and ubiquitinated proteins were immunoprecipitated from whole-cell lysates (WCL) (IP: anti-HA). Immunoblots of IPs and WCL were probed for RIP2 (anti-FLAG) and ubiquitin (anti-HA). Data are representative of three independent experiments. **(B)** K27-linked polyubiquitin chains suppress NOD2 signaling. HEK293 cells co-transfected with plasmids encoding NOD2, FLAG-RIP2, an NF-κB luciferase reporter, and the indicated HA-tagged ubiquitin constructs were stimulated or not with L18-MDP for 60 min, followed by analysis of NF-κB luciferase activity. Data are shown as mean ± SEM (n=3 independent experiments, each in triplicate). Statistical analysis was conducted using one-way ANOVA followed by Tukey’s multiple comparisons test. *p* < 0.05; p < 0.01; *p* < 0.001. **(C)** K27-linked polyubiquitin chains suppress NOD2-driven cytokine responses. ELISA quantification of IL-6 and TNF-α secretion from HEK293 cells transfected and stimulated as in (B) for 24 h. Data are shown as mean ± SEM (n=3 independent experiments). Statistical significance was determined using one-way ANOVA with Tukey’s post hoc test. *p* < 0.05; p < 0.01. **(D)** HEK293 cells co-transfected with plasmids encoding NOD2, FLAG-RIP2, and HA-tagged ubiquitin constructs (WT, KO, K6-only, K11-only, K27-only, K33-only, K48-only, K63-only, or K63+K27-only) were stimulated with L18-MDP for 0 or 60 min. FLAG-immunoprecipitated RIP2 complexes and whole-cell lysates (WCLs) were immunoblotted for the indicated signaling components (TAK1, p-TAK1, IKKα/β, p-IKKα/β, TAB2, TAB3, NEMO). **(E)** Schematic model illustrating the effects of different ubiquitin linkage types on RIP2 signaling. K63- and M1-linked polyubiquitin chains promote NOD2-driven inflammatory signaling, whereas K27-linked chains suppress RIP2-mediated responses, highlighting their potential role in negative regulation.

To determine how K27-linked chains block NOD2-RIP2 signaling, we examined their impact on RIP2 signalosome formation. RIP2 activation recruits TAK1, TAB2/3, and the IKK complex (IKKα/β/NEMO) to drive inflammation. HEK293 cells co-transfected with FLAG-RIP2, NOD2, and HA-tagged ubiquitin constructs (WT, K63-only, K27-only, K63+K27) were stimulated with L18-MDP, followed by RIP2 immunoprecipitation. HA-K63 Ub enhanced recruitment of TAK1, TAB2/3, and NEMO to RIP2, with increased p-TAK1 and p-IKKα/β (Fig 2D). In contrast, HA-K27 Ub strongly reduced recruitment of these effectors and lowered their phosphorylation in lysates. The K63+K27 variant showed intermediate impairment versus K63 alone, confirming K27’s dominant-negative effect (Fig. 2D). Together, these data demonstrate opposing roles of ubiquitin linkages in NOD2–RIP2 signaling, whereby K63 and M1 linkages promote pro-inflammatory responses, whereas K27 linkages antagonize signaling by disrupting RIP2 signalosome assembly and suppressing NOD2-driven inflammatory signaling (Fig 2E).

### XIAP Catalyzes Dual K63/K27 RIP2 Ubiquitination to Fine-Tune NOD2 Signaling

To identify the E3 ligase mediating RIP2 K27 ubiquitination, we examined XIAP (X-linked inhibitor of apoptosis protein) - a known component of the RIP2 complex that catalyzes K63-RIP2 ubiquitination and scaffolds LUBAC in NOD2 signaling (Damgaard *et al*., 2012; Goncharov *et al*., 2018; Krieg *et al*., 2009; Stafford *et al*., 2018).

In WT BMDMs treated with Smac mimetic LCL161 (IAP inhibitor) and stimulated with L18-MDP (0-60 min), RIP2 immunoprecipitation revealed rapid, parallel K63/K27 induction peaking at 30-60 min. LCL161 selectively abolished K27 but not K63-ubiquitination, implicating XIAP in K27-chain formation (Fig 3A).

**Figure 3.**
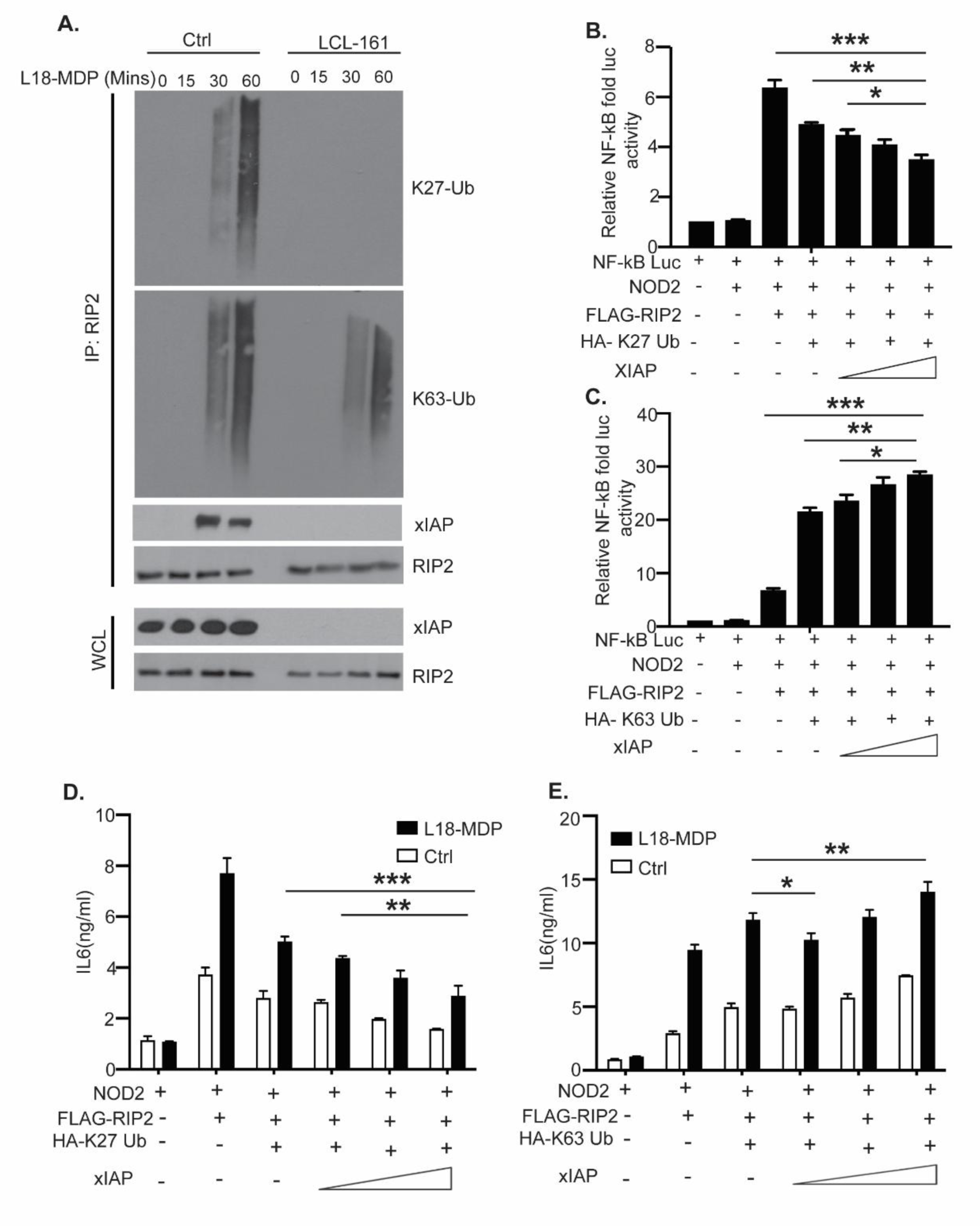
XIAP bidirectionally regulates NOD2-RIP2 signaling via K27- and K63-linked ubiquitination. (A) Inhibition of XIAP abolishes NOD2-induced K27-linked polyubiquitination of RIP2. Wild-type BMDMs stimulated with L18-MDP (0, 15, 30, 60 min) ± LCL161 (Smac mimetic) were subjected to RIP2 immunoprecipitation and immunoblotting for K27- and K63-linked ubiquitin. (B) XIAP suppresses NOD2 signaling via K27-linked polyubiquitination. NF-κB luciferase activity in HEK293 cells co-transfected with NOD2, FLAG-RIP2, NF-κB reporter, HA-K27 Ub, and increasing XIAP amounts. (C) XIAP enhances NOD2 signaling via K63-linked polyubiquitination. As in (B), but with HA-K63 Ub. Data are representative of 2-3 independent experiments. (**D, E**) XIAP bidirectionally regulates NOD2-driven cytokine secretion via K27 and K63-linked polyubiquitination. ELISA quantification of IL-6 from HEK293 cells co-transfected with NOD2, FLAG-RIP2, HA-K27 Ub (D) or HA-K63 Ub (E) and increasing XIAP amounts.

XIAP overexpression with HA-K63 Ub enhanced NF-κB-luciferase activity (Fig 3B), whereas HA-K27 Ub overexpression caused dose-dependent suppression (Fig 3C). Cytokine ELISAs (IL-6/TNFα) mirrored this pattern: K63 increased output; K27 reduced it (Fig 3D and E; Fig EV1A and B).

Integrated with evidence of LUBAC-mediated M1 extensions on both chains on RIP2 (Panda & Gekara, 2018), these results position XIAP as a versatile E3 ligase initiating activating (K63) and repressive (K27) signals on RIP2, enabling multilayered ubiquitin control of NOD2 signaling as a molecular rheostat.

These findings reveal a complex ubiquitin landscape on RIP2, where XIAP acts as a versatile E3 ligase - driving signal propagation via K63-linked chains while counterbalancing it through repressive K27-linked modifications - establishing combinatorial ubiquitination as a rheostat for NOD2 signaling.

### XIAP E3 Ligase Activity Drives K27 RIP2 Ubiquitination and NOD2 Signaling Suppression

To assess whether the E3 ligase activity of XIAP is required to suppress NOD2-RIP2 signaling, we used a catalytically inactive mutant (XIAP-H467A) carrying a histidine-to-alanine substitution at position 467 in the RING domain (Gyrd-Hansen *et al*, 2008). HEK293 cells were co-transfected with NOD2, RIP2, HA-K27 Ub, and either WT XIAP or the catalytically inactive XIAP-H467A mutant. In the presence of HA-K27 Ub, WT XIAP markedly suppressed NOD2-induced NF-κB luciferase activity (Fig EV2A) and reduced cytokine release (IL-6/TNFα) post-stimulation (Fig EV 2B and C). In contrast, the H467A mutant failed to inhibit signaling, confirming XIAP E3 ligase activity is essential for downstream suppression. These results demonstrate XIAP limits NOD2-RIP2-driven NF-κB activation and cytokine production in an E3-dependent manner, consistent with inhibitory K27 ubiquitination of RIP2 to restrain inflammation (Fig EV2D).

### MYSM1 Counteracts XIAP by Removing Inhibitory K27 Chains from RIP2

E3 ligases like XIAP often pair with deubiquitinases (DUBs) to dynamically regulate signaling. MYSM1 was previously shown to suppress RIP2 activity by cleaving activating K63/M1 chains (Panda & Gekara, 2018). However, its capacity for K27 cleavage remained untested in this context.

To test MYSM1’s removal of inhibitory K27 chains from RIP2, HEK293 cells were co-transfected with an NF-κB luciferase reporter, NOD2, FLAG-RIP2, and HA-K27 Ub or HA-K63 Ub together with increasing doses of MYSM1. HA-K27 Ub strongly suppressed NOD2-induced NF-κB activation, which MYSM1 reversed dose-dependently (Fig 4B). Conversely, HA-K63 Ub enhanced activation, an effect that was attenuated by MYSM1 (Fig 4C). Cytokine ELISAs (IL-6/TNFα) mirrored these trends without altering RIP2 levels, confirming direct chain editing (Fig 4D and E; Fig EV3). MYSM1 relieved K27-mediated inhibition of IL-6 and TNFα secretion and reduced the K63-driven overproduction of these cytokines. These effects occurred without changes in RIP2 protein abundance, indicating that MYSM1 acts directly on ubiquitin chains rather than indirectly by modulating RIP2 complex assembly or stability. Collectively, these findings identify MYSM1 as a critical regulator of NOD2–RIP2 signaling that fine-tunes pathway output by controlling both inhibitory (K27-linked) and activating (K63-linked) ubiquitination events on RIP2 (Fig 4E).

**Figure 4.**
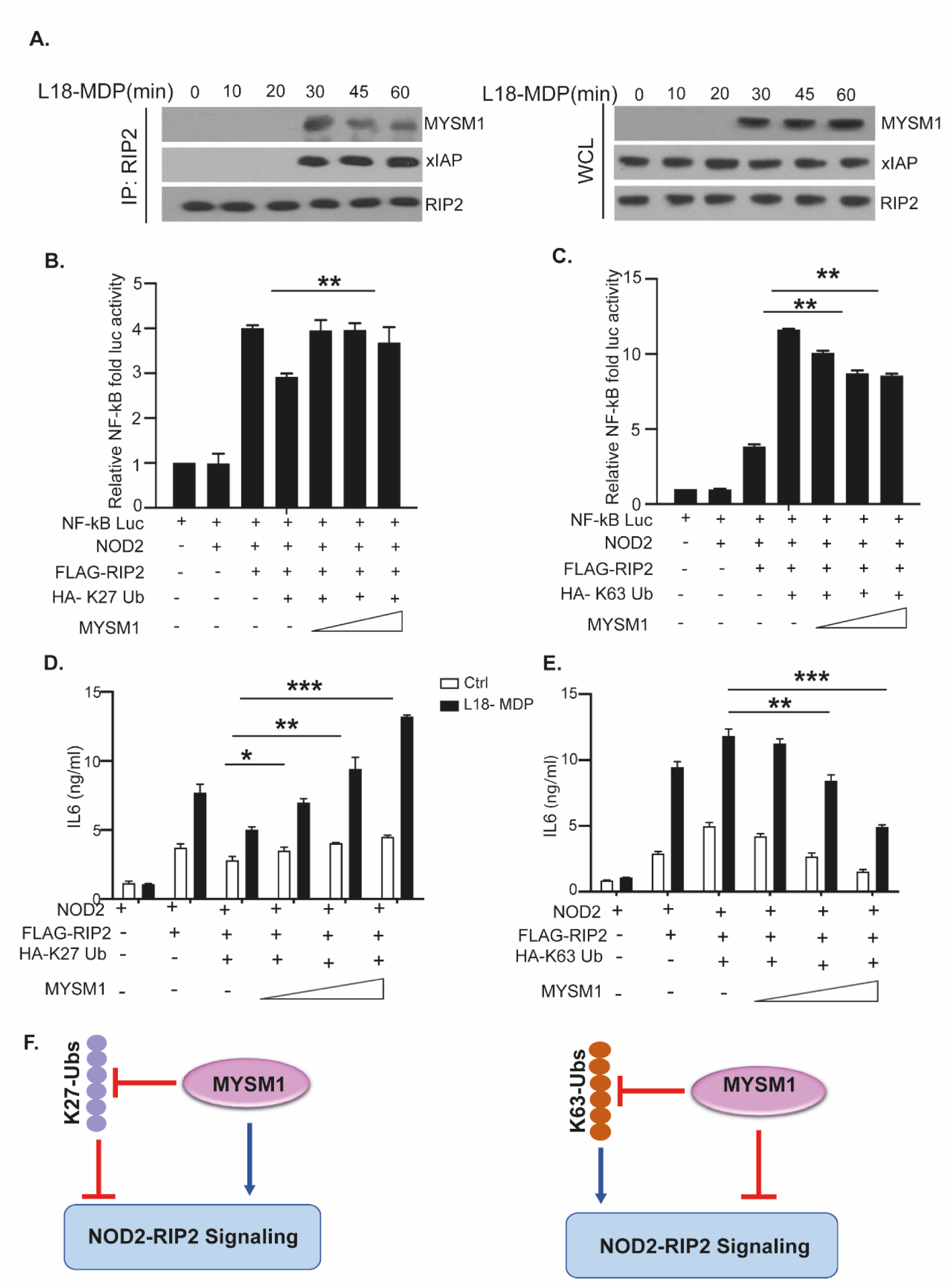
MYSM1 controls K27- and K63-linked ubiquitination of RIP2 to bidirectionally regulate NOD2 signaling. (A) MYSM1 associates with the RIP2–XIAP signaling complex. RIP2 was immunoprecipitated from wild-type BMDMs stimulated with L18-MDP for the indicated times, and immunoprecipitates were immunoblotted for MYSM1 and XIAP. (B) MYSM1 enhances NOD2 signaling by promoting K27-linked polyubiquitination of RIP2. NF-κB luciferase activity was measured in HEK293 cells co-transfected with NOD2, FLAG–RIP2, an NF-κB reporter construct, HA–K27-only ubiquitin, and increasing amounts of MYSM1, then stimulated with L18-MDP for 60 min or left unstimulated. (C) MYSM1 attenuates NOD2 signaling by regulating K63-linked polyubiquitination of RIP2. NF-κB luciferase activity was measured in HEK293 cells co-transfected with NOD2, FLAG–RIP2, an NF-κB reporter, HA–K63-only ubiquitin, and increasing amounts of MYSM1, then stimulated with L18-MDP (60 min) or left unstimulated. Data represent the mean ± SEM of three independent experiments performed in triplicate. Statistical significance was determined by one-way ANOVA with Tukey’s post hoc test (*p < 0.05; **p < 0.01; **p < 0.001). (D) MYSM1 promotes NOD2-driven cytokine production by regulating K27-linked polyubiquitination of RIP2. IL-6 secretion was quantified by ELISA in HEK293 cells co-transfected with NOD2, FLAG–RIP2, HA–K27-only ubiquitin, and increasing amounts of MYSM1, then stimulated with L18-MDP (60 min) or left unstimulated. (E) MYSM1 restrains NOD2-driven cytokine production by regulating K63-linked polyubiquitination of RIP2. IL-6 levels were measured by ELISA in HEK293 cells co-transfected with NOD2, FLAG–RIP2, HA–K63-only ubiquitin, and increasing amounts of MYSM1, then stimulated with L18-MDP (60 min) or left unstimulated. Data represent the mean ± SEM of three independent experiments. Statistical significance was determined by one-way ANOVA with Tukey’s multiple comparisons test (*p < 0.05; *p < 0.01). (F) Schematic model illustrating MYSM1 as a dual-function deubiquitinase that removes both activating (K63-linked) and inhibitory (K27-linked) ubiquitin chains from RIP2, thereby fine-tuning NOD2-induced inflammatory signaling.

### MYSM1 controls NOD2-RIP2 signaling via its SWIRM and MPN domains

MYSM1 harbors two conserved domains critical for its activity: the SWIRM domain, which mediates protein–protein and chromatin interactions, and the MPN (Mpr1/Pad1 N-terminal) domain, which provides catalytic deubiquitinase function(Panda & Gekara, 2018; Panda *et al*, 2015). To define the minimal MYSM1 elements required for regulation of NOD2–RIP2 signaling, we generated a recombinant fusion protein containing only the SWIRM and MPN domains (SWIRM+MPN) and assessed its activity in an *in vitro* deubiquitinase assay. In L18-MDP-stimulated WT BMDMs, immunoprecipitated RIP2 carried K63-, K48-, K27-, and M1-linked polyubiquitin chains. Incubation with SWIRM+MPN selectively reduced K63, K27, and M1 linkages while sparing K48 chains (Fig 5A).

**Figure 5.**
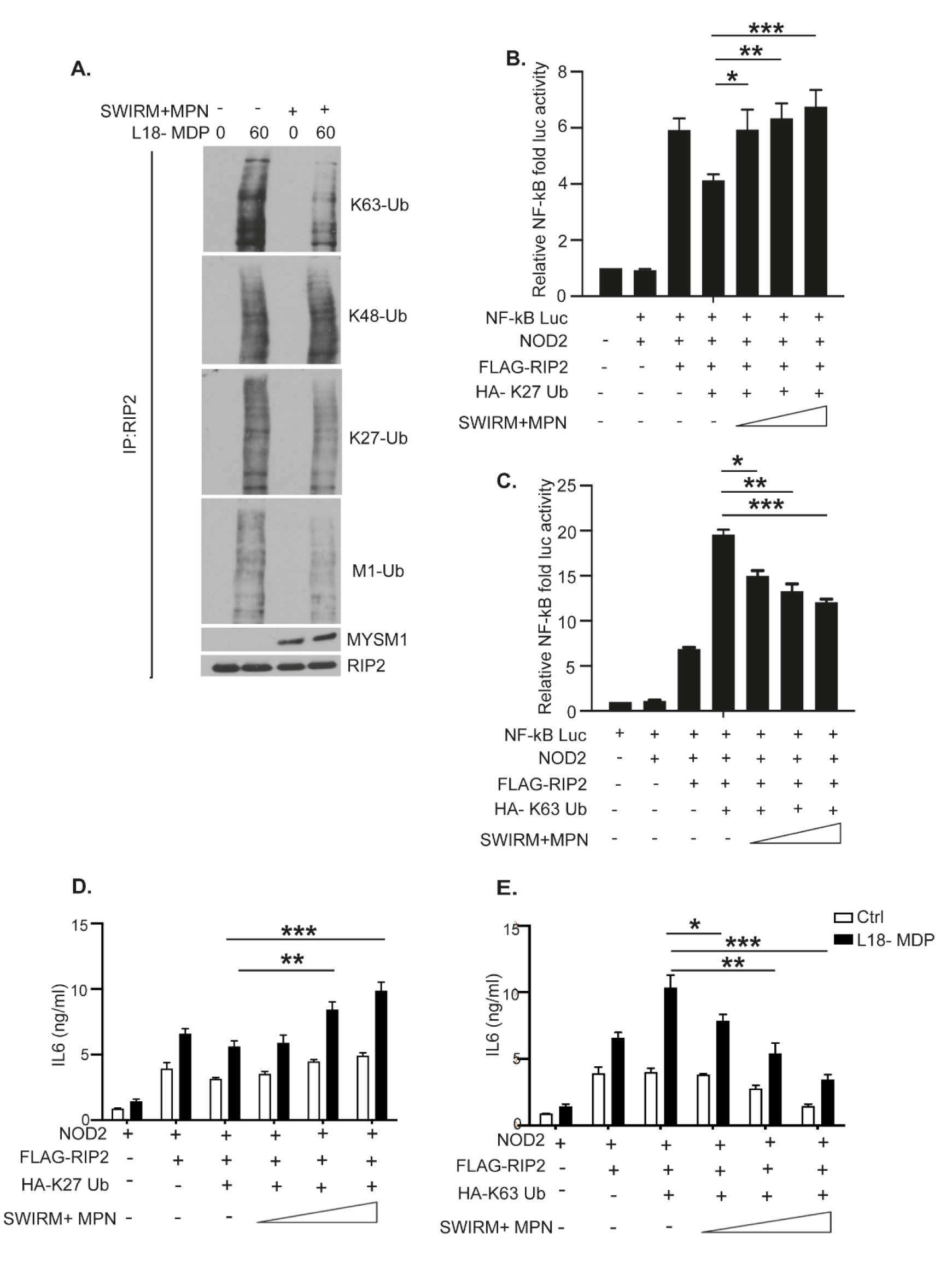
The SWIRM and MPN domains of MYSM1 regulate NOD2–RIP2 signaling through selective targeting of K63- and K27-linked ubiquitin chains. **(A)** MYSM1’s SWIRM+MPN domain edit K27 and K63-linked polyubiquitins from RIP2. RIP2 immunoprecipitated from wild type BMDMs stimulated (or not) with L18-MDP for indicated duration, was incubated with recombinant fusion of SWIRM and MPN domains on MYSM1 (SWIRM+MPN). The IPs were then immunoblotted with antibodies against the indicated linkage-specific antibodies. **(B)** MYSM1’s SWIRM+MPN domain enhances NOD2 signaling by editing K27-linked polyubiquitination from RIP2. NF-κB luciferase activity in HEK293 cells that were co-transfected with NOD2, FLAG-RIP2, an NF-κB reporter, HA-K27-only ubiquitin, and increasing amounts of MYSM1, then stimulated with L18-MDP (60 min) or left unstimulated. Data represent mean ± SEM from three independent experiments performed in triplicate. Statistical significance was determined using one-way ANOVA with Tukey’s multiple comparisons test. *p* < 0.05; p < 0.01; *p* < 0.001. **(C)** MYSM1’s SWIRM+MPN domain restrains NOD2 signaling by editing K63-linked polyubiquitination from RIP2. NF-κB luciferase activity in HEK293 cells that were co-transfected with NOD2, FLAG–RIP2, an NF-κB reporter, HA-K63-only ubiquitin, and increasing amounts of MYSM1, then stimulated with L18-MDP (60 min) or left unstimulated. Data represent mean ± SEM from three independent experiments performed in triplicate. Statistical significance was determined using one-way ANOVA with Tukey’s multiple comparisons test. *p* < 0.05; p < 0.01; *p* < 0.001. **(D)** MYSM1’s SWIRM+MPN domain promotes NOD2-driven cytokine production by regulating K27-linked polyubiquitination of RIP2. ELISA quantification of IL-6 secretion by HEK293 cells that were co-transfected with NOD2, FLAG-RIP2, HA-K27-only ubiquitin, and increasing amounts of MYSM1 then stimulated with L18-MDP (60 min) or left unstimulated. **(E)** MYSM1 restrains NOD2-driven cytokine production by editing K63-linked polyubiquitination from RIP2. ELISA quantification of IL-6 secretion by HEK293 cells that were co-transfected with NOD2, FLAG-RIP2, HA-K63-only ubiquitin, and increasing amounts of MYSM1, then stimulatied with L18-MDP (60 min) or left unstimulated. Results are shown as mean ± SEM from three independent experiments. Statistical analysis was conducted using one-way ANOVA followed by Tukey’s post hoc test. *p* < 0.05; p < 0.01.

Next, we next overexpressed the SWIRM+MPN construct alongside NF-κB luciferase reporter plasmids, NOD2, FLAG-RIP2, and either K27- or K63-linked ubiquitin. The SWIRM+MPN restored NF-κB luciferase activity suppressed by K27-linked ubiquitin and reduced the hyperactivation induced by K63-linked chains (Fig 5B and C). These effects mirrored those observed with full-length MYSM1, demonstrating that the dual-domain module alone is sufficient for linkage-specific deubiquitination of RIP2 (Fig4). Consistently, analysis of IL-6 and TNFα secretion revealed that SWIRM+MPN expression could reverse K27-mediated suppression and normalized the elevated cytokine output driven by K63 linkages (Fig 5D and E; Fig EV4).

Collectively, these results establish that the SWIRM and MPN domains form a self-contained, catalytically active unit capable of regulating RIP2 signaling by selectively editing both inhibitory (K27-linked) and activating (K63-linked) polyubiquitin chains.

## Discussion

In this study we identify K27-linked polyubiquitination of RIP2 emerges as a negative regulatory node in the NOD2 signaling pathway. Whereas the activating roles of K63- and M1-linked polyubiquitin chains in promoting RIP2 complex assembly and driving NF-κB and MAPK activation are well established (Damgaard *et al*., 2012; Jiang *et al*, 2024; Panda & Gekara, 2018; Tao *et al*., 2009; Tokunaga *et al*., 2009), our data show that K27 linkages act in direct opposition, functioning as an intrinsic brake that limits RIP2 signalosome formation and inflammatory output.

Mechanistically, we demonstrate that K27-linked chains impair the recruitment of key adaptors including TAK1, TAB2/3, and NEMO to RIP2, likely via steric or conformational constraints that disrupt protein-protein interactions or alter RIP2 oligomerization. In mixed-linkage architectures containing K63 chains, K27 linkages dominantly suppress signaling, attenuating the assembly or stability of activating scaffolds (Fig.2).

We identify XIAP as the E3 ligase catalyzing K27-linked ubiquitination of RIP2. While XIAP has been implicated in promoting RIP2 activation through K63- and M1-linked chains(Damgaard *et al*., 2012; Jiang *et al*., 2024; Panda & Gekara, 2018; Tao *et al*., 2009; Tokunaga *et al*., 2009), our findings reveal a bifunctional role: in addition to activating chains, XIAP deposits inhibitory K27 linkages. Pharmacological inhibition of XIAP with the Smac mimetic LC161 selectively abolishes K27, but not K63, ubiquitination, indicating mechanistic separation of these activities. Consistently, the catalytically inactive H467A XIAP mutant fails to impose K27-linked repression of NOD2–RIP2 signaling (Fig EV2). Overexpression of K27-linked ubiquitin alone is sufficient to dampen NF-κB activity and cytokine production, even under upstream NOD2 activation (Fig 2).

These findings redefine XIAP as a dual-function ligase that concurrently installs activating (K63) and inhibitory (K27) chains, enabling dynamic tuning of signal amplitude. We propose a model in which XIAP rapidly conjugates both linkage types early in the response, establishing a built-in negative feedback circuit to constrain activation. LUBAC-mediated M1 extensions may subsequently stabilize these mixed-linkage scaffolds, embedding additional layers of control. The interplay of K63, K27, and M1 thus encodes a combinatorial ubiquitin logic that operates as a molecular rheostat to calibrate immune output.

Acting downstream, MYSM1 serves as a broad-spectrum counter-regulatory deubiquitinase that edits both activating (K63 and M1) chains as well as inhibitory (K27) chains from RIP2. Unlike more linkage-restricted DUBs such as A20, CYLD, or OTULIN (Fiil *et al*, 2013; Hitotsumatsu *et al*, 2008; Hrdinka *et al*., 2016; Panda & Gekara, 2018; Witt & Vucic, 2017), MYSM1 removes either chain type, reversing K27-mediated repression and tempering K63 and M1-driven hyperactivation. We define a minimal SWIRM-MPN module sufficient for this dual activity, restoring signaling from K27 repression and normalizing cytokine output under K63-driven stimulation (Fig 4 and 5; Fig EV3 and EV4).

Together, these findings define the XIAP-RIP2-MYSM1 axis as a tunable signaling module in which immune signaling output is governed by the dynamic balance between selective ubiquitin assembly and context-dependent removal. K27-linked chains act as reversible inhibitory modifications on RIP2, serving to restrain excessive activation. XIAP installs both activating and inhibitory ubiquitin linkages, whereas MYSM1 removes both types to reset the signaling state. This coordinated interplay ensures that immune responses remain robust yet self-limiting, promoting effective host defense while preventing inflammatory overactivation.

Dysregulated NOD2-RIP2 signaling underlies chronic inflammatory diseases, including Crohn’s disease (Hugot *et al*., 2001) and Blau syndrome (Mao *et al*., 2022). The discovery of K27-linked ubiquitination as an inhibitory ubiquitin code, dynamically regulated by the XIAP-MYSM1 axis, defines a tunable checkpoint in this pathway and a promising therapeutic target. MYSM1’s unique linkage-specific editing capacity further enables precise immunomodulation to reinstate immune homeostasis.

## Methods

### Reagents and Tools Table

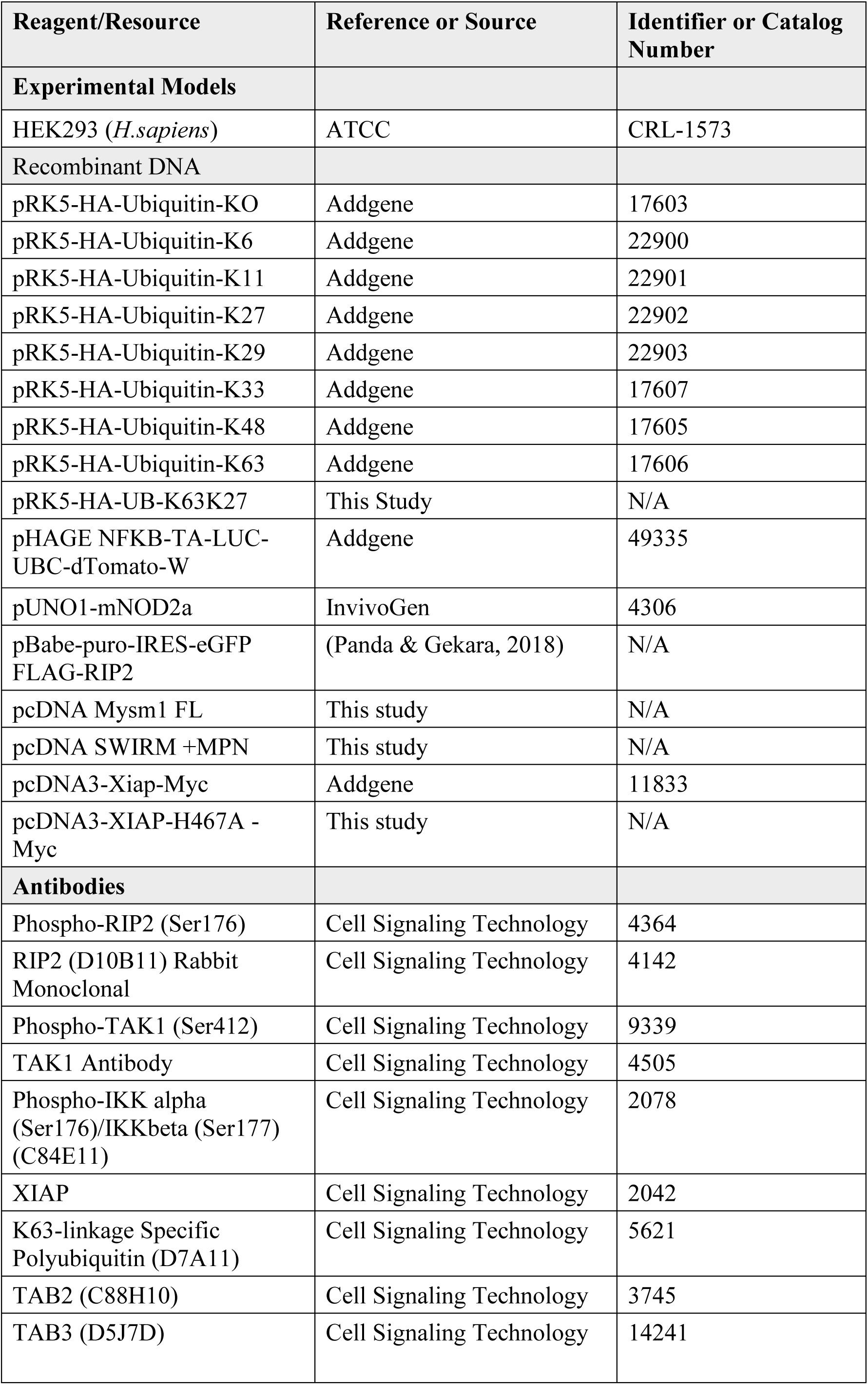
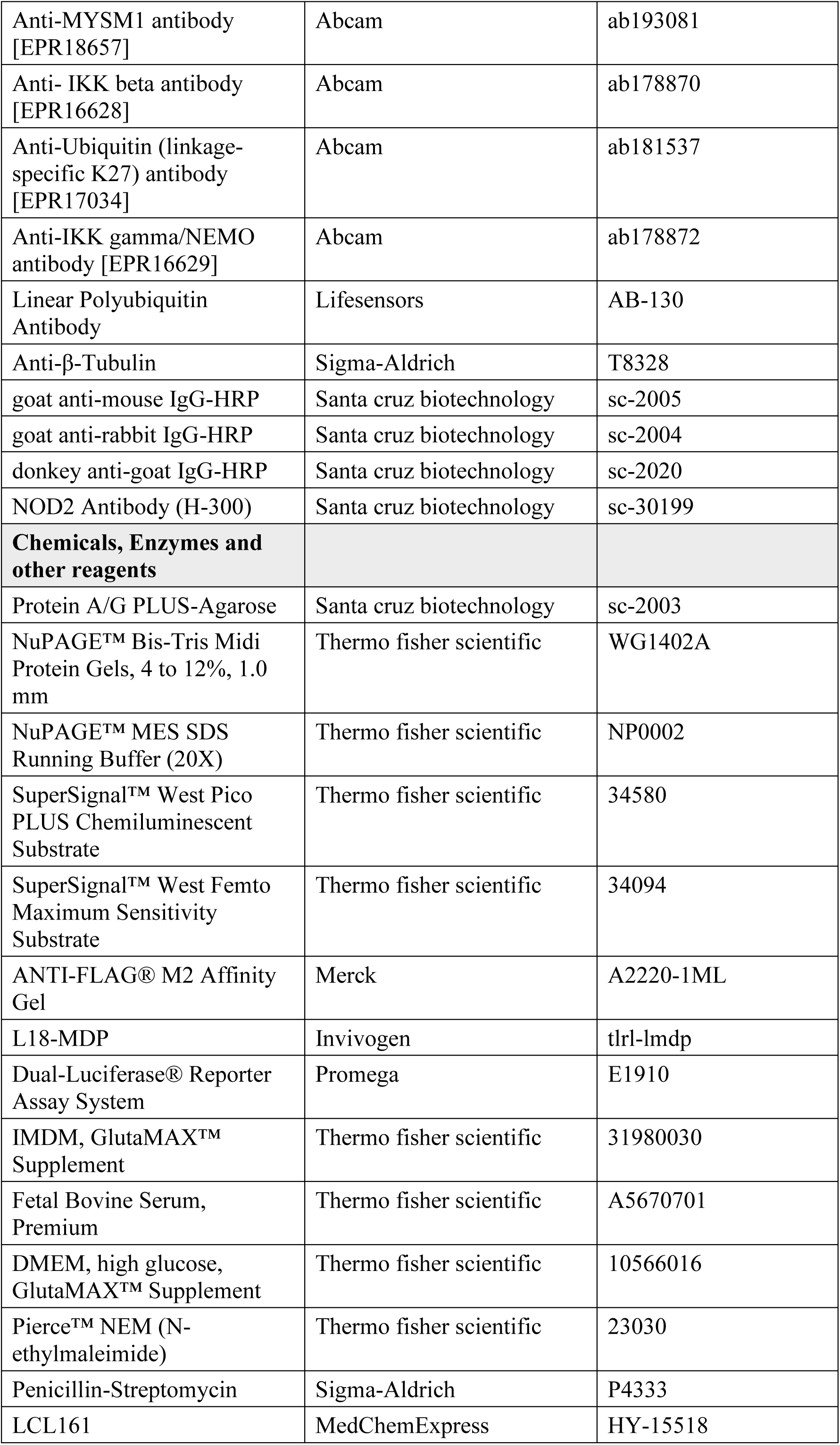
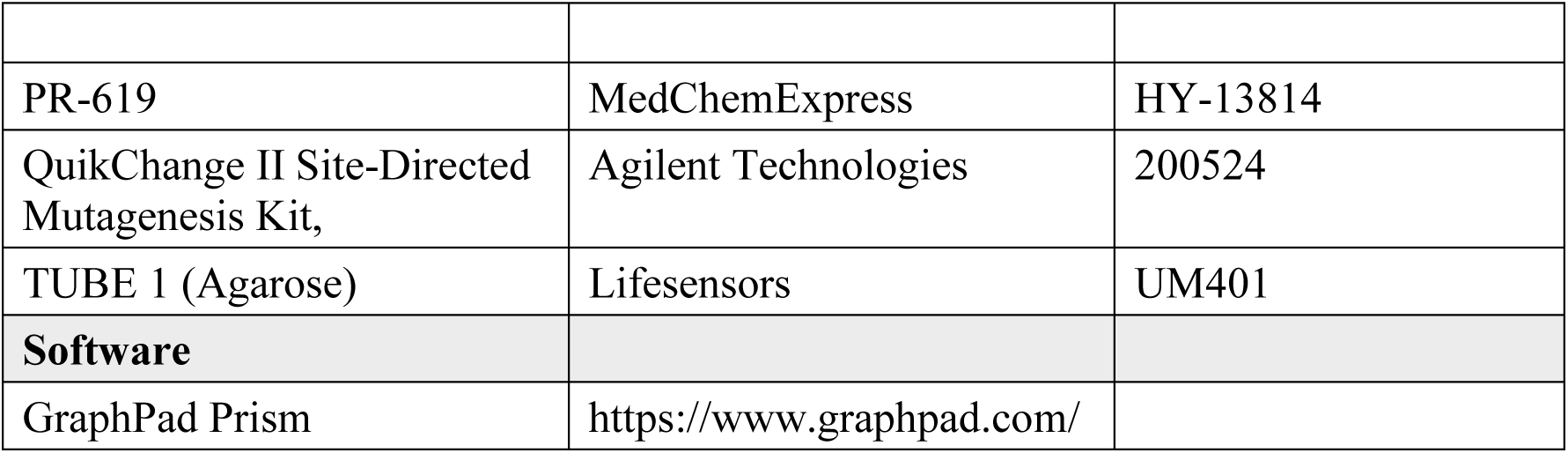

## Ethics statement

All animal were maintained in accordance with institutional guidelines and approved by the Swedish Board of Agriculture (Jordbruks verket; approval no. A53-14) under specific pathogen-free (SPF) conditions.

### Mice

Breeding of mice was done at the Umeå Transgene Facility (UTCF).

### Plasmids and Constructs

Full-length FLAG-tagged RIP2 was cloned into the pBabe-puro-IRES-EGFP vector (Addgene plasmid #14430). HA-tagged ubiquitin variants-wild-type, K0, K27-only, K63-only, and k63K27-only were expressed from pRK5 vectors (Addgene). The pUNO1-NOD2 construct was purchased from InvivoGen, and pcDNA3-XIAP-Myc was obtained from Addgene. The XIAP-H467A catalytic mutant was generated by site-directed mutagenesis using the QuikChange II kit (Agilent Technologies), verified by Sanger sequencing, and subcloned into pcDNA3.1. Full-length MYSM1 cDNA and its SWIRM + MPN catalytic domain fusion were cloned into pcDNA3.1 for mammalian expression. All cloned constructs were sequence-verified prior to use.

### Cell culture and stimulation

BMDMs were generated were generated by culturing bone marrow progenitors in IMDM (1×) +Glutamax™-I (Gibco, Life technology) supplemented with 10% FCS (Gibco, Life technology), 100 U/ml penicillin/streptomycin (Sigma-Aldrich) and 20% (v/v) L929 conditional medium for 5 days. HEK293 cells (ATCC) were maintained in DMEM (1×) +Glutamax™-I (Gibco, Life technology) supplemented with 10% FCS (Gibco, Life technology) and 100 U/ml penicillin/streptomycin (Sigma-Aldrich). For induction of inflammatory signaling, BMDMs were stimulated by adding L18-MDP (200 ng/ml; InvivoGen) directly to the culture medium. For inhibitor studies, BMDMs were pre-treated with (or without) SMAC mimetic (a IAPs inhibitor, LCL-161) (100 nM) for 18h prior to L18-MDP stimulation.

### Transient Transfection and Reporter Assays

HEK293 cells were seeded in 24-well plates and transfected using Lipofectamine 2000 (Thermo fisher) with the indicated expression plasmids. After 48 h, cells were stimulated with L18-MDP (1µg/mL) for 60 min and harvested for luciferase measurement using the Dual-Luciferase Reporter Assay System (Promega). Firefly luciferase activity was normalized to Renilla luciferase. Data are presented as mean ± SEM of at least three independent experiments performed in triplicate.

### Immunoprecipitation and immunoblot analysis

For standard immunoblotting, cells were lysed directly in 2× Laemmli buffer and analyzed by SDS-PAGE followed by immunoblotting. For immunoprecipitation (IP) assays, cells were lysed in buffer containing 1% NP-40, 50 mM Tris-HCl (pH 7.5), 150 mM NaCl, 1 mM NaF, 2 mM PMSF, 1 mM sodium orthovanadate, 10 mM sodium pyrophosphate, and a protease inhibitor cocktail (Roche Applied Science). For denaturing IPs, lysates were resuspended in 100 μl of TSD buffer (50 mM Tris-HCl [pH 7.5], 1% SDS, 5 mM DTT) and boiled for 10 min. After centrifugation (13,000 rpm, 5 min, room temperature), supernatants were diluted in 1.2 ml of TNN buffer (50 mM Tris-HCl [pH 7.5], 250 mM NaCl, 5 mM EDTA, 0.5% NP-40). Samples were incubated with specific antibodies overnight at 4 °C with gentle rotation, and immune complexes were captured using Protein A/G agarose beads. Beads were washed three times with lysis buffer, resuspended in Laemmli buffer, and analyzed by SDS-PAGE and immunoblotting.

For ubiquitination assays, cells were lysed in urea lysis buffer (8 M urea, 50 mM Tris-HCl [pH 7.5], 25 mM NaCl, 5 mM EDTA, 2 mM N-ethylmaleimide, and protease inhibitors). Lysates were sonicated and centrifuged (14,000 rpm, 10 min) to remove debris. Supernatants were mixed with reducing SDS sample buffer, boiled at 70 °C for 10 min, and subjected to SDS-PAGE followed by immunoblotting with anti-ubiquitin antibodies.

### Purification of endogenous total ubiquitin conjugates

Endogenous ubiquitin (Ub) conjugates were purified using tandem ubiquitin-binding entities (TUBEs; LifeSensors). Cells were lysed in TUBE lysis buffer (50 mM Tris-HCl [pH 7.5], 150 mM NaCl, 1 mM EDTA, 1% NP-40, 10% glycerol) supplemented with 5 mM N-ethylmaleimide (NEM), 20 μM PR-619 (pan-deubiquitinase inhibitor), and a protease inhibitor cocktail (Roche). Lysates were cleared by centrifugation. For TUBE pulldowns, agarose-conjugated TUBEs were equilibrated according to the manufacturer’s instructions (LifeSensors) and incubated with clarified lysates at 4 °C overnight with gentle agitation. Beads were washed three times with ice-cold TUBE lysis buffer and resuspended in Laemmli buffer for SDS-PAGE and immunoblot analysis.

### Expression and purification of recombinant MYSM1

The mouse Mysm1 gene encoding a His-SWIRM-MPN fusion was cloned into the *pET30a* expression vector and transformed into E. coli *BL21 (DE3)* cells. Bacterial cultures (1 L) were grown in M9 medium at 37 °C until mid-log phase and protein expression was induced with 1 mM IPTG at 15 °C overnight. Cells were harvested by centrifugation, resuspended in lysis buffer (50 mM Tris-HCl [pH 8.0], supplemented with PMSF), and lysed by sonication. After centrifugation to remove cell debris, the supernatant was subjected to nickel-affinity chromatography. Bound His-SWIRM-MPN fusion protein was eluted using a stepwise imidazole gradient (Panda & Gekara, 2018; Panda *et al*., 2015).

### In vitro deubiquitination assay

For in vitro deubiquitination assays, ubiquitin-conjugated protein complexes bound to beads were resuspended in 1× deubiquitinase (DUB) buffer (125 mM HEPES [pH 7.5], 25 mM MgCl₂, 10 mM NaF, 10 nM okadaic acid, 500 mM NaCl, 2.5 mM DTT, and 0.5 mg/ml BSA). Recombinant his-SWIRM+MPN (10 μl; 1 μg) was added to the bead slurry and incubated for 30 min at 37 °C with gentle agitation. The reaction was terminated by adding 2× SDS loading buffer, and samples were analyzed by SDS-PAGE followed by immunoblotting using anti-K63-, anti-K48-, anti-K27-, and anti-M1-linked ubiquitin antibodies.

### Cytokine Measurements by ELISA

Cell-free supernatants were collected from transfected or stimulated cells, centrifuged, and assayed for IL-6 and TNF-α using ELISA kits (R&D Systems) according to the manufacturer’s protocol. Absorbance was read at 450 nm. Cytokine concentrations were determined from standard curves and expressed as mean ± SEM of ≥ 3 biological replicates.

## Statistical Analysis

All data are presented as mean ± SEM from at least three independent biological replicates. Statistical analyses were performed using GraphPad Prism 10. Unless otherwise indicated, comparisons between groups were analyzed using one-way ANOVA followed by Bonferroni’s multiple comparison post-test. *P-values < 0.05, **P < 0.01, ***P < 0.001; were considered to be significant.

## Data availability

All data supporting the conclusions of this study are included in the main text and/or the Expanded View (EV) files.

## Acknowledgements

This work was funded by Stockholm University, The Laboratory for Molecular Infection Medicine Sweden (MIMS) - Umea University, The MBW - Stockholm University – Sweden and The Medical Center - University of Freiburg to N.O.G. The Swedish Research Council (2016-00890 and 2022-01308_3 to N.O.G), The Swedish Cancer Foundation (CAN 20 1252PjF01H, CAN 23 3096 Pj, to N.O.G.

## Author contributions

S.P and N.O.G conceived the study and designed the experiments. SP and AK performed the experiments.

S.P and N.O.G analyzed and interpreted the data. NOG supervised the research along with S.P. S.P and

N.O.G wrote the manuscript, with comments from AK.

## Expanded View Figures

**Figure EV1.**
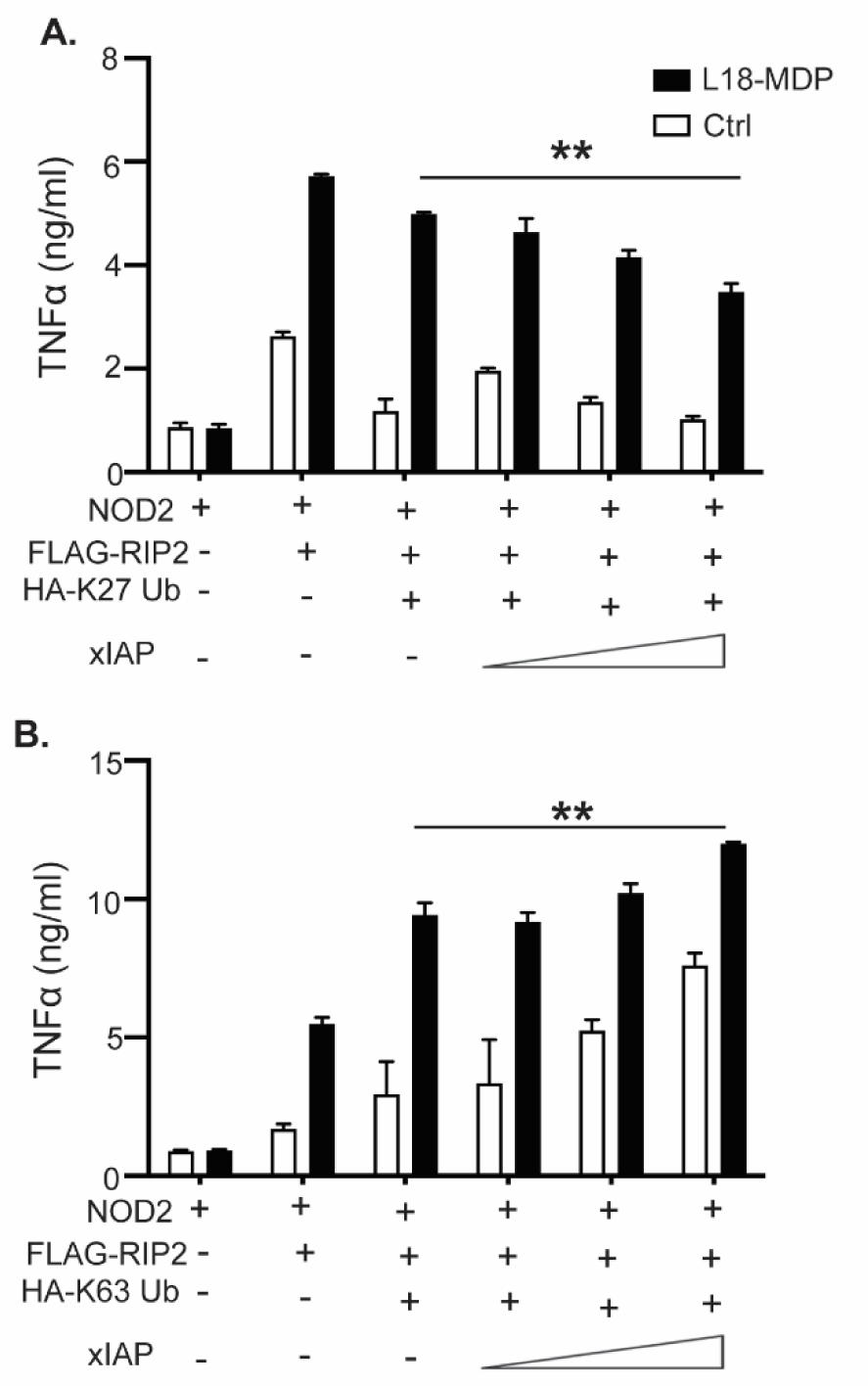
XIAP bidirectionally regulates the NOD2-driven cytokine response through K27- and K63-linked ubiquitination. (A, **B)** XIAP bidirectionally regulates NOD2-driven cytokine secretion via K27 and K63-linked polyubiquitination. ELISA quantification of TNF-α from HEK293 cells that were co-transfected with NOD2, FLAG-RIP2, HA-K27 Ub (A) or HA-K63 Ub (B), and increasing XIAP amounts, then stimulated with L18-MDP or left unstimulated. Data are representative of three independent experiments.

**Figure EV2.**
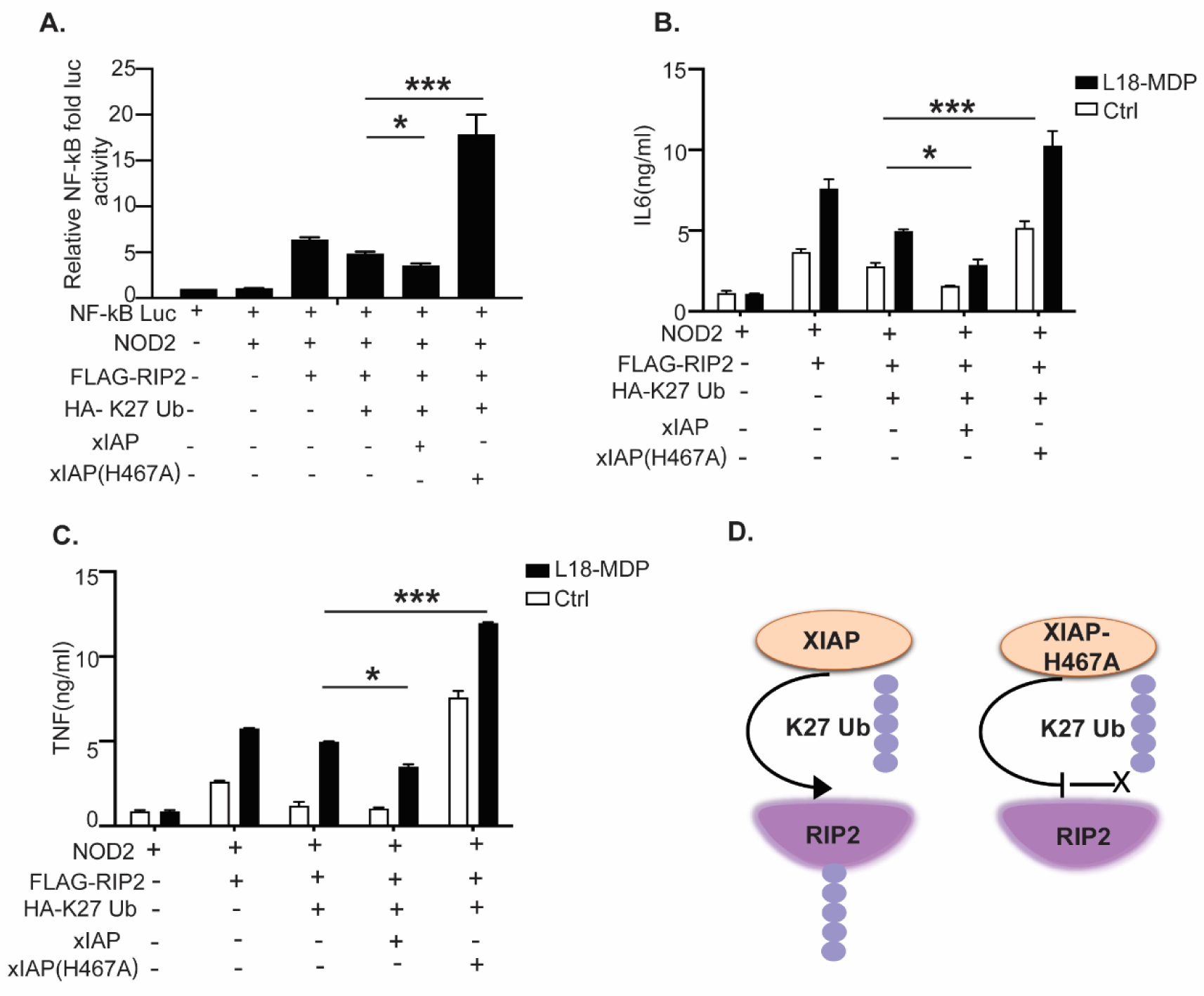
The E3 ligase activity of XIAP is required to suppress NOD2–RIP2 signaling through K27-linked ubiquitination. **(A)** NF-κB luciferase reporter assay in HEK293 cells co-transfected with NOD2, RIP2, K27-only ubiquitin, and either wild-type XIAP or the catalytically inactive H467A mutant. **(B, C)** ELISA quantification of IL-6 (B) TNFα (C) secretion by HEK293 cells co-transfected with plasmids for NOD2, RIP2, K27-only ubiquitin and either wild-type or H467A XIAP. **(D)** Schematic depicting XIAP-mediated ubiquitination of RIP2 with K27-linked chains, which suppress downstream NF-κB activation. The catalytically inactive H467A mutant fails to conjugate K27-linked ubiquitin to RIP2. Data are representative of at least two independent experiments; error bars denote ±SEM.

**Figure EV3.**
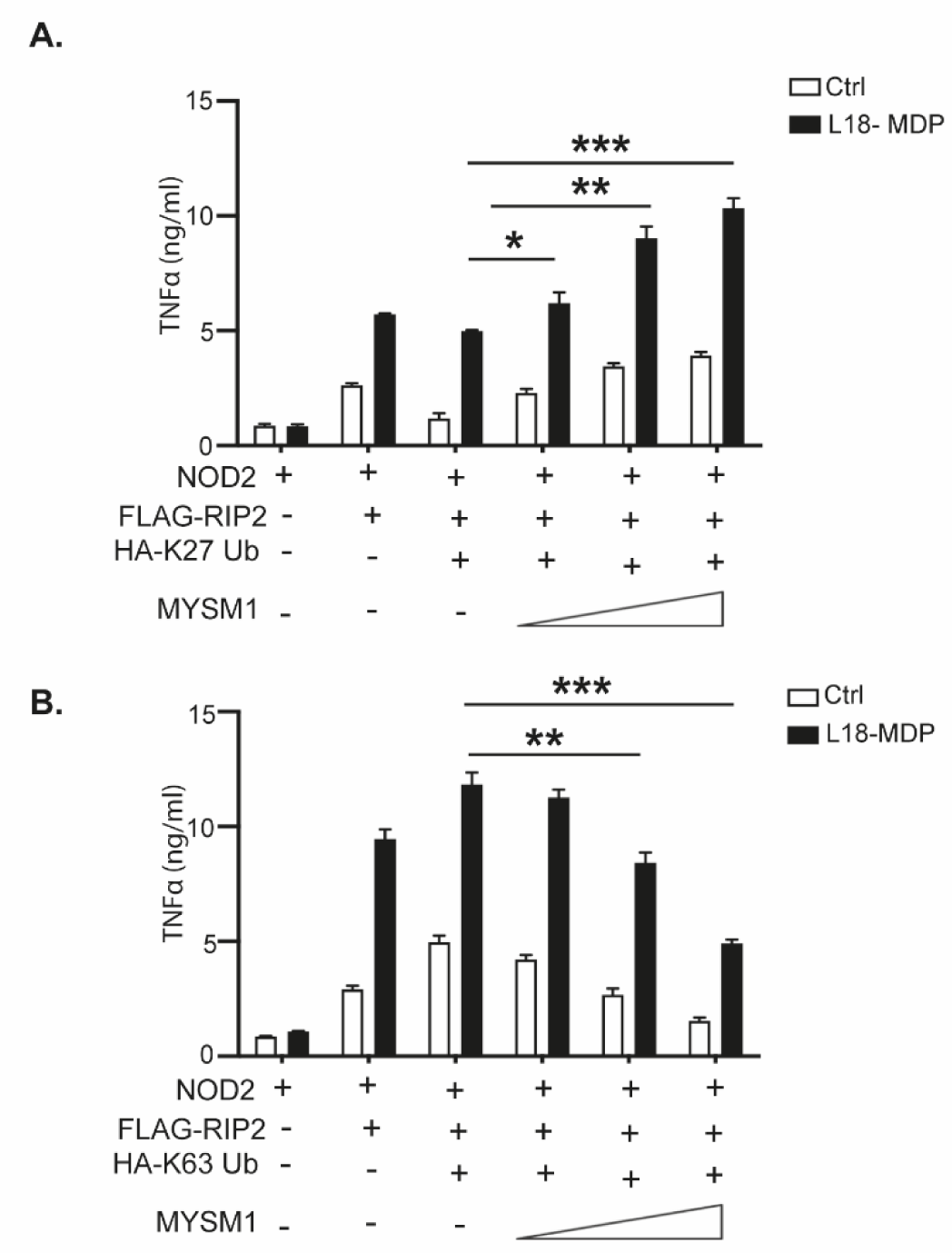
MYSM1 bidirectionally regulates NOD2–RIP2–driven cytokine responses through K27- and K63-linked polyubiquitination. (A, **B)** ELISA quantification of TNFα secretion in HEK293 cells that were co-transfected with plasmids expressing NOD2, FLAG–RIP2, HA–K27-only ubiquitin (A) or HA–K63-only ubiquitin (B), and increasing amounts of MYSM1, followed by stimulation with L18-MDP or left unstimulated.

**Supplementary Figure 4.**
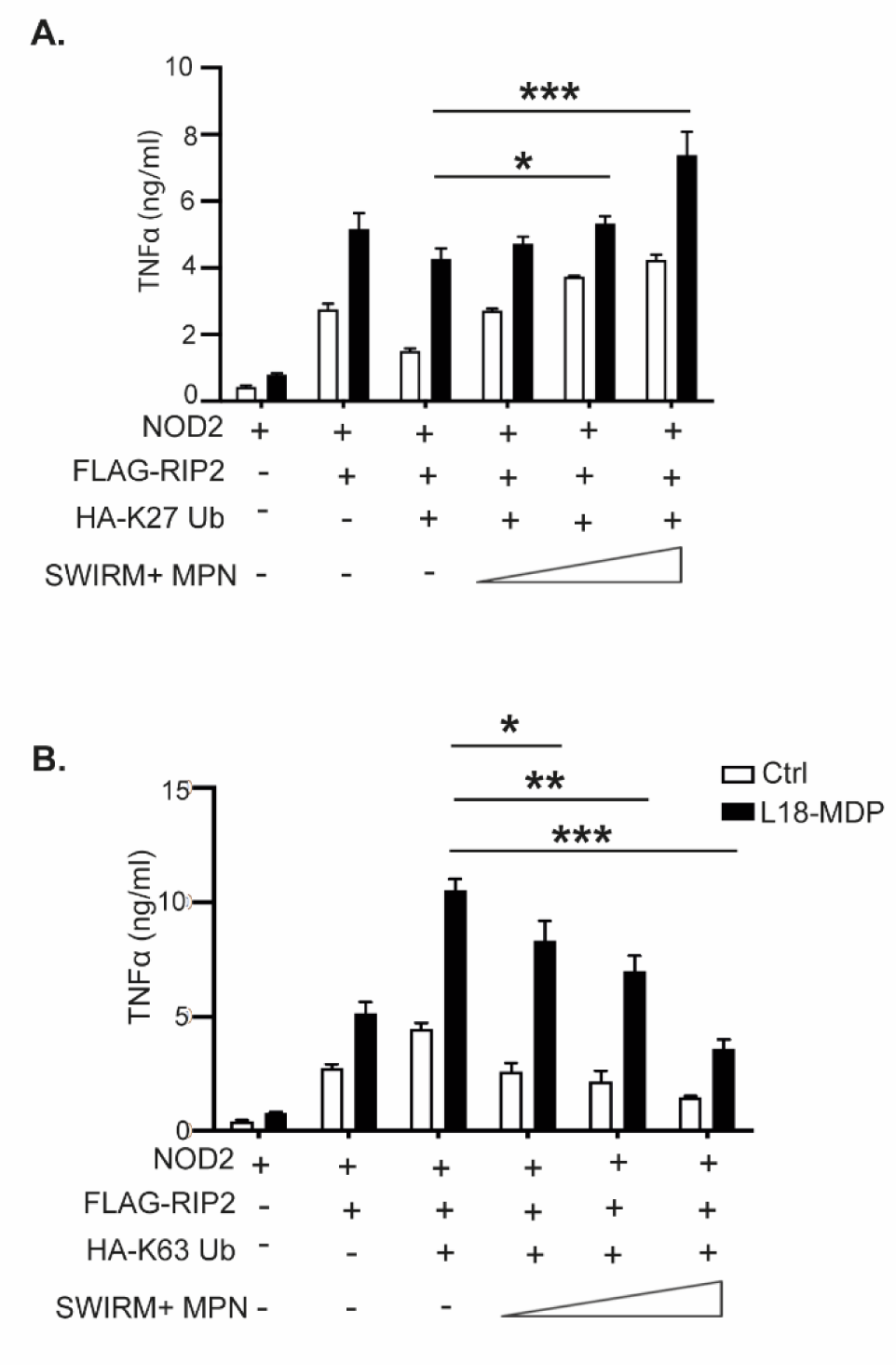
The SWIRM and MPN domains of MYSM1 regulate NOD2–RIP2-driven cytokine response by targeting K63 and K27-llnked polyubiquitins. (A, **B)** ELISA quantification of TNFα secretion in HEK293 cells that were with plasmids encoding NOD2, Flag-RIP2, HA-K27-only (A) or K63-only (B) Ubs and increasing amounts of SWIRM+MPN, then stimulated (or not) with L18-MDP. Data represent mean ± SEM from three independent experiments. Statistical significance was determined using one-way ANOVA followed by Tukey’s post hoc test. *p* < 0.05; **p** < 0.01.

## References

1. Agrata R, Komander D (2025) Ubiquitin-A structural perspective. Mol Cell 85: 323–346

2. Akira S, Uematsu S, Takeuchi O (2006) Pathogen recognition and innate immunity. Cell 124: 783–801

3. Akizuki Y, Kaypee S, Ohtake F, Ikeda F (2024) The emerging roles of non-canonical ubiquitination in proteostasis and beyond. J Cell Biol 223

4. Akutsu M, Dikic I, Bremm A (2016) Ubiquitin chain diversity at a glance. J Cell Sci 129: 875–880

5. Bertrand MJ, Doiron K, Labbé K, Korneluk RG, Barker PA, Saleh M (2009) Cellular inhibitors of apoptosis cIAP1 and cIAP2 are required for innate immunity signaling by the pattern recognition receptors NOD1 and NOD2. Immunity 30: 789–801

6. Bhoj VG, Chen ZJ (2009) Ubiquitylation in innate and adaptive immunity. Nature 458: 430–437

7. Castañeda CA, Dixon EK, Walker O, Chaturvedi A, Nakasone MA, Curtis JE, Reed MR, Krueger S, Cropp TA, Fushman D (2016) Linkage via K27 Bestows Ubiquitin Chains with Unique Properties among Polyubiquitins. Structure 24: 423–436

8. Chau V, Tobias JW, Bachmair A, Marriott D, Ecker DJ, Gonda DK, Varshavsky A (1989) A multiubiquitin chain is confined to specific lysine in a targeted short-lived protein. Science 243: 1576–1583

9. Chen ZJ, Sun LJ (2009) Nonproteolytic functions of ubiquitin in cell signaling. Mol Cell 33: 275–286

10. Damgaard RB (2021) The ubiquitin system: from cell signalling to disease biology and new therapeutic opportunities. Cell Death Differ 28: 423–426

11. Damgaard RB, Nachbur U, Yabal M, Wong WW, Fiil BK, Kastirr M, Rieser E, Rickard JA, Bankovacki A, Peschel C et al (2012) The ubiquitin ligase XIAP recruits LUBAC for NOD2 signaling in inflammation and innate immunity. Mol Cell 46: 746–758

12. Draber P, Kupka S, Reichert M, Draberova H, Lafont E, de Miguel D, Spilgies L, Surinova S, Taraborrelli L, Hartwig T et al (2015) LUBAC-Recruited CYLD and A20 Regulate Gene Activation and Cell Death by Exerting Opposing Effects on Linear Ubiquitin in Signaling Complexes. Cell Rep 13: 2258–2272

13. Fiil BK, Damgaard RB, Wagner SA, Keusekotten K, Fritsch M, Bekker-Jensen S, Mailand N, Choudhary C, Komander D, Gyrd-Hansen M (2013) OTULIN restricts Met1-linked ubiquitination to control innate immune signaling. Mol Cell 50: 818–830

14. Gatti M, Pinato S, Maiolica A, Rocchio F, Prato MG, Aebersold R, Penengo L (2015) RNF168 promotes noncanonical K27 ubiquitination to signal DNA damage. Cell Rep 10: 226–238

15. Girardin SE, Boneca IG, Viala J, Chamaillard M, Labigne A, Thomas G, Philpott DJ, Sansonetti PJ (2003) Nod2 is a general sensor of peptidoglycan through muramyl dipeptide (MDP) detection. J Biol Chem 278: 8869–8872

16. Goncharov T, Hedayati S, Mulvihill MM, Izrael-Tomasevic A, Zobel K, Jeet S, Fedorova AV, Eidenschenk C, deVoss J, Yu K et al (2018) Disruption of XIAP-RIP2 Association Blocks NOD2-Mediated Inflammatory Signaling. Mol Cell 69: 551–565.e557

17. Gyrd-Hansen M, Darding M, Miasari M, Santoro MM, Zender L, Xue W, Tenev T, da Fonseca PC, Zvelebil M, Bujnicki JM et al (2008) IAPs contain an evolutionarily conserved ubiquitin-binding domain that regulates NF-kappaB as well as cell survival and oncogenesis. Nat Cell Biol 10: 1309–1317

18. Harris LD, Le Pen J, Scholz N, Mieszczanek J, Vaughan N, Davis S, Berridge G, Kessler BM, Bienz M, Licchesi JDF (2021) The deubiquitinase TRABID stabilizes the K29/K48-specific E3 ubiquitin ligase HECTD1. J Biol Chem 296: 100246

19. Hitotsumatsu O, Ahmad RC, Tavares R, Wang M, Philpott D, Turer EE, Lee BL, Shiffin N, Advincula R, Malynn BA et al (2008) The ubiquitin-editing enzyme A20 restricts nucleotide-binding oligomerization domain containing 2-triggered signals. Immunity 28: 381–390

20. Hrdinka M, Fiil BK, Zucca M, Leske D, Bagola K, Yabal M, Elliott PR, Damgaard RB, Komander D, Jost PJ et al (2016) CYLD Limits Lys63- and Met1-Linked Ubiquitin at Receptor Complexes to Regulate Innate Immune Signaling. Cell Rep 14: 2846–2858

21. Hu H, Sun SC (2016) Ubiquitin signaling in immune responses. Cell Res 26: 457–483

22. Hugot JP, Chamaillard M, Zouali H, Lesage S, Cézard JP, Belaiche J, Almer S, Tysk C, O’Morain CA, Gassull M et al (2001) Association of NOD2 leucine-rich repeat variants with susceptibility to Crohn’s disease. Nature 411: 599–603

23. Jiang W, Li X, Xu H, Gu X, Li S, Zhu L, Lu J, Duan X, Li W, Fang M (2023) UBL7 enhances antiviral innate immunity by promoting Lys27-linked polyubiquitination of MAVS. Cell Rep 42: 112272

24. Jiang W, Zhao Y, Han M, Xu J, Chen K, Liang Y, Yin J, Hu J, Shen Y (2024) N4BP3 facilitates NOD2-MAPK/NF-κB pathway in inflammatory bowel disease through mediating K63-linked RIPK2 ubiquitination. Cell Death Discov 10: 440

25. Komander D, Reyes-Turcu F, Licchesi JD, Odenwaelder P, Wilkinson KD, Barford D (2009) Molecular discrimination of structurally equivalent Lys 63-linked and linear polyubiquitin chains. EMBO Rep 10: 466–473

26. Krieg A, Correa RG, Garrison JB, Le Negrate G, Welsh K, Huang Z, Knoefel WT, Reed JC (2009) XIAP mediates NOD signaling via interaction with RIP2. Proc Natl Acad Sci U S A 106: 14524–14529

27. Li J, Chai QY, Liu CH (2016) The ubiquitin system: a critical regulator of innate immunity and pathogen-host interactions. Cell Mol Immunol 13: 560–576

28. Liu J, Qian C, Cao X (2016) Post-Translational Modification Control of Innate Immunity. Immunity 45: 15–30

29. Liu W, Wang Y, Liu S, Zhang X, Cao X, Jiang M (2024) E3 Ubiquitin Ligase RNF13 Suppresses TLR Lysosomal Degradation by Promoting LAMP-1 Proteasomal Degradation. Adv Sci (Weinh) 11: e2309560

30. Lu C, Liu H, Liu T, Sun S, Zheng Y, Ling T, Luo X, E Y, Xu Y, Li J et al (2025) RIPK2 promotes colorectal cancer metastasis by protecting YAP degradation from ITCH-mediated ubiquitination. Cell Death Dis 16: 248

31. Mao L, Dhar A, Meng G, Fuss I, Montgomery-Recht K, Yang Z, Xu Q, Kitani A, Strober W (2022) Blau syndrome NOD2 mutations result in loss of NOD2 cross-regulatory function. Front Immunol 13: 988862

32. Michel MA, Swatek KN, Hospenthal MK, Komander D (2017) Ubiquitin Linkage-Specific Affimers Reveal Insights into K6-Linked Ubiquitin Signaling. Mol Cell 68: 233–246.e235

33. Panda S, Gekara NO (2018) The deubiquitinase MYSM1 dampens NOD2-mediated inflammation and tissue damage by inactivating the RIP2 complex. Nat Commun 9: 4654

34. Panda S, Nilsson JA, Gekara NO (2015) Deubiquitinase MYSM1 Regulates Innate Immunity through Inactivation of TRAF3 and TRAF6 Complexes. Immunity 43: 647–659

35. Parackova Z, Bloomfield M, Vrabcova P, Zentsova I, Klocperk A, Milota T, Svaton M, Casanova JL, Bustamante J, Fronkova E et al (2020) Mutual alteration of NOD2-associated Blau syndrome and IFNγR1 deficiency. J Clin Immunol 40: 165–178

36. Park JH, Kim YG, McDonald C, Kanneganti TD, Hasegawa M, Body-Malapel M, Inohara N, Núñez G (2007) RICK/RIP2 mediates innate immune responses induced through Nod1 and Nod2 but not TLRs. J Immunol 178: 2380–2386

37. Pellegrini E, Desfosses A, Wallmann A, Schulze WM, Rehbein K, Mas P, Signor L, Gaudon S, Zenkeviciute G, Hons M et al (2018) RIP2 filament formation is required for NOD2 dependent NF-κB signalling. Nat Commun 9: 4043

38. Shearer RF, Typas D, Coscia F, Schovsbo S, Kruse T, Mund A, Mailand N (2022) K27-linked ubiquitylation promotes p97 substrate processing and is essential for cell proliferation. EMBO J 41: e110145

39. Sheng X, Xia Z, Yang H, Hu R (2024) The ubiquitin codes in cellular stress responses. Protein Cell 15: 157–190

40. Stafford CA, Lawlor KE, Heim VJ, Bankovacki A, Bernardini JP, Silke J, Nachbur U (2018) IAPs Regulate Distinct Innate Immune Pathways to Co-ordinate the Response to Bacterial Peptidoglycans. Cell Rep 22: 1496–1508

41. Takeuchi O, Akira S (2010) Pattern recognition receptors and inflammation. Cell 140: 805–820 Tao M, Scacheri PC, Marinis JM, Harhaj EW, Matesic LE, Abbott DW (2009) ITCH K63-ubiquitinates the NOD2 binding protein, RIP2, to influence inflammatory signaling pathways. *Curr Biol* 19: 1255-1263

42. Tokunaga F, Sakata S, Saeki Y, Satomi Y, Kirisako T, Kamei K, Nakagawa T, Kato M, Murata S, Yamaoka S et al (2009) Involvement of linear polyubiquitylation of NEMO in NF-kappaB activation. Nat Cell Biol 11: 123–132

43. Windheim M, Lang C, Peggie M, Plater LA, Cohen P (2007) Molecular mechanisms involved in the regulation of cytokine production by muramyl dipeptide. Biochem J 404: 179–190

44. Witt A, Vucic D (2017) Diverse ubiquitin linkages regulate RIP kinases-mediated inflammatory and cell death signaling. Cell Death Differ 24: 1160–1171

45. Yang S, Wang B, Humphries F, Jackson R, Healy ME, Bergin R, Aviello G, Hall B, McNamara D, Darby T et al (2013) Pellino3 ubiquitinates RIP2 and mediates Nod2-induced signaling and protective effects in colitis. Nat Immunol 14: 927–936

46. Yau R, Rape M (2016) The increasing complexity of the ubiquitin code. Nat Cell Biol 18: 579–586 Zhang J, Luo Y, Wu B, Huang X, Zhao M, Wu N, Miao J, Li J, Zhu L, Wu D et al (2024) Identifying functional dysregulation of NOD2 variant Q902K in patients with Yao syndrome. *Arthritis Res Ther* 26: 58

47. Zhao J, Cai B, Shao Z, Zhang L, Zheng Y, Ma C, Yi F, Liu B, Gao C (2021) TRIM26 positively regulates the inflammatory immune response through K11-linked ubiquitination of TAB1. Cell Death Differ 28: 3077–3091

